# Heat-inactivated mycobacteria activate the Toll-like receptor 2 and 4 pathways in the zebrafish model of tuberculosis

**DOI:** 10.1101/2023.03.31.535083

**Authors:** Elisa Ferreras-Colino, Marinela Contreras, María A. Risalde, Iker A. Sevilla, Encarnación Delgado, Lucas Domínguez, Christian Gortazar, Jose de la Fuente

**Affiliations:** SaBio, Instituto de Investigación en Recursos Cinegéticos IREC-CSIC-UCLM-JCCM, Ronda de Toledo 12, 13005 Ciudad Real, Spain; Department of Veterinary Pathobiology, Center for Veterinary Health Sciences, Oklahoma State University, Stillwater, OK 74078, USA; Departamento de Anatomía y Anatomía Patológica Comparadas y Toxicología, Grupo de Investigación GISAZ, UIC Zoonosis y Enfermedades Emergentes ENZOEM, Universidad de Córdoba, Córdoba, Spain; CIBERINFEC, ISCIII - CIBER de Enfermedades Infecciosas, Instituto de Salud Carlos III, Madrid, Spain; Animal Health Department, NEIKER – Basque Institute for Agricultural Research and Development, Basque Research and Technology Alliance (BRTA), 48160 Derio, Bizkaia, Spain; VISAVET Health Surveillance Center. Universidad Complutense de Madrid, Madrid, Spain

**Keywords:** fish mycobacteriosis, inactivated *Mycobacterium bovis*, Toll-like receptors

## Abstract

Based on previous evidence demonstrating the efficacy of inactivated mycobacteria for the control of fish mycobacteriosis, we explored the protective efficacy of two inactivated *Mycobacterium bovis* administered via parenteral and mucosal routes against *Mycobacterium marinum* infection mimicking natural conditions in the zebrafish model of tuberculosis. Although we did not observe a clear effect of any of the immunostimulants on mycobacterial burden, the results showed a significant increase in TLR2 and TLR4 gene expression levels in fishes parenterally immunized with inactivated Bacillus Calmette-Guérin (BCG). Our findings demonstrated that the TLR2 and the TLR4 signaling pathways are involved in the immune response elicited by inactivated mycobacteria in the zebrafish model of tuberculosis and support the use of inactivated mycobacteria in vaccine formulations for the control of mycobacteriosis.

## 1. Introduction

Fish mycobacteriosis is a chronic disease caused mainly by *Mycobacterium marinum* (*M. marinum*) that affects fresh and marine water fish, including wild, aquaculture and ornamental species [1–3]. Due to the economic detriment for the aquaculture industry, as well as the zoonotic risk [1–3], cost-effective measures are required for the control of fish mycobacteriosis. Since mycobacterial infections generally require antibiotherapy for long periods of time, together with the growing spread of antibiotic resistance, immunoprophylaxis arises as the strategy of choice for effective and sustainable disease control. Yet, no licensed vaccines against fish mycobacteriosis are currently commercialized [1,4]. Numerous approaches aiming to induce a protective immune response against *Mycobacterium* spp. in fish have been attempted (reviewed in [1]), several of them achieving promising effects. In particular, vaccination with inactivated *M. marinum* [4] or with the related mycobacterium *M. bovis*, either alive (BCG) [5] or heat-inactivated (HIMB) [2,3], has demonstrated protection against fish mycobacteriosis, partly attributed to the stimulation of the innate immune system.

In the last few decades, the zebrafish (*Danio rerio*) has arisen as a validated animal model for the research of mycobacterial infections [6–8]. The versatility of the species allows evaluating diverse immunization and infection routes [7], such as the parenteral administration via intraperitoneal injection [3] of the mucosal administration via bath immersion [2], therefore providing a wide range of possibilities to transfer into the aquaculture field.

The objective of the present study was to compare the protective efficacy of two inactivated*M. bovis*, iBCG and HIMB, as well as the administration route, parenteral[3] or mucosal[2], against *M. marinum* simulating natural infection (sublethal dose via mucosal route) in the zebrafish model of tuberculosis.

## 2. Material and Methods

### 2.1. Ethics Statement

Zebrafish were provided by Juan Galcerán Sáez from the Instituto de Neurociencias (IN-CSIC-UMH, Sant Joan d’Alacant, Alicante, Spain) and were certified by Biosait Europe S.L. (Barcelona, Spain; https://biosait.com) as free of major fish pathogens including *Mycobacterium* spp. Animals were maintained in a flow-through water system at 27 °C with a light/dark cycle of 14 h/10 h and fed twice daily with dry fish feed (Premium food tropical fish, DAPC, Valladolid, Spain). Experiments in zebrafish were conducted in strict accordance with the recommendations of the European Guide for the Care and Use of Laboratory Animals. Fish were housed and experiments conducted at experimental facility (IREC, Ciudad Real, Spain) with the approval and supervision of the Ethics Committee on Animal Experimentation of the University of Castilla La Mancha (PR-2021-09-14) and the Counseling of Agriculture, Environment and Rural Development of Castilla La Mancha (REGA code ES130340000218).

### 2.2. Animals and experimental design

One hundred twenty adult (6–8 months old) wild-type AB female and male zebrafish were randomly assigned to five groups with intraperitoneal (IP) or immersion (IMM) immunostimulant administration. **Group 1** received two doses of 20 μl of 10^4^ colony forming units (CFU) of HIMB each via IP inoculation with an interval of 3 weeks; **Group 2** received two doses of 20 μl of 10^4^ CFU of iBCG each via IP inoculation with an interval of 3 weeks; **Group 3** was immersed in 400 ml of 10^5^ CFU of HIMB for 30 minutes with an interval of 3 weeks; **Group 4** was immersed in 400 ml of 10^5^ CFU of iBCG for 30 minutes with an interval of 3 weeks; **Group 5** received PBS instead (Figure 1). Two weeks after the last immunization, all animals were infected via immersion with approximately 10^4^ *M. marinum* CFU per fish in 400 ml of water for 30 minutes. Three weeks after infection, all animals were euthanized with tetracayne [2,3] and were sampled for molecular studies or processed for histopathological studies.

**Figure 1.**
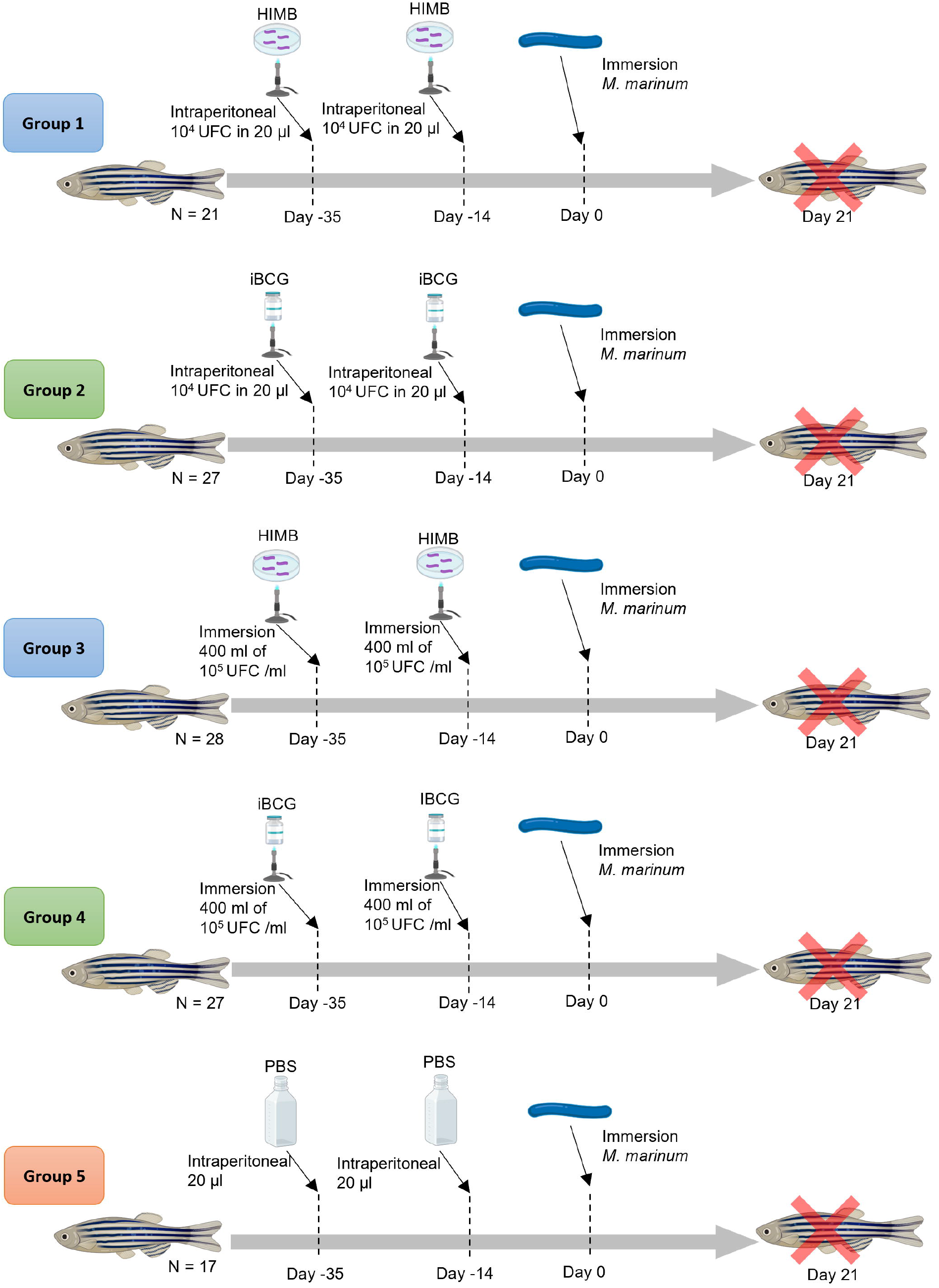
Experimental design. One hundred twenty adult (6–8 months old) wild-type AB female and male zebrafish were randomly assigned to five groups **Group 1** received two doses of 20 μl of 10^4^ colony forming units (CFU) of HIMB each via IP inoculation with an interval of 3 weeks; **Group 2** received two doses of 20 μl of 10^4^ CFU of iBCG each via IP inoculation with an interval of 3 weeks; **Group 3** was immersed in 400 ml of 10^5^ CFU of HIMB with an interval of 3 weeks; **Group 4** was immersed in 400 ml of 10^5^ CFU of iBCG with an interval of 3 weeks; **Group 5** received PBS instead (Figure 1). Fourteen days after the last immunization, all animals were infected via immersion with approximately 10^4^ *M. marinum* CFU per fish in 400 ml of water.

### 2.3. Immunostimulants

#### Heat-inactivated <u>M. bovis</u>

HIMB immunostimulant consisted of 20 μl of sterile PBS containing approximately 10^4^ CFU/ml for parenteral administration or 400 ml of water containing approximately 10^5^ CFU/ml for mucosal administration of a field isolate (Strain 1403; SB0339 spoligotype) obtained from a naturally infected wild boar. HIMB was prepared as described in Vaz-Rodrigues et al. [9].

#### *Inactivated* <u>Bacillus Calmette-Guérin</u>

Inactivated BCG immunostimulant consisted of 20 μl of sterile PBS containing approximately 10^4^ CFU/ml for parenteral administration or 400 ml of water containing approximately 10^5^ CFU/ml for mucosal administration of BCG Danish. BCG was inactivated as described for HIMB in Vaz-Rodrigues et al. [9].

### 2.4. M. marinum

*M. marinum* (ATCC 927) was cultured at 29 °C in 7H9 broth enriched with Middlebrook ADC (Becton Dickinson) and prepared for infection as was previously described [2,3] to adjust to an approximate dose of 10^4^ CFU of *M. marinum* per fish.

### 2.5. Quantitative polymerase chain reaction (qPCR)

Total DNA and mRNA were isolated from the intestine samples collected during fish necropsy with the AllPrep DNA/RNA/Protein Kit (Qiagen, Hilden, Germany), following manufacturer’s instructions. Nucleic acids concentration and absorbance ratio at 260/280 nm were assessed with the Nanodrop One spectrophotometer (Thermo Scientific, Wilmington, DE, USA) at an optical density of 260 nm (OD260).

Relative gene expression of *M. marinum* 16S rDNA and *D. rerio* immune mediators were assessed by qPCR with the SsoFast™ EvaGreen^®^ Supermix (BioRad, Hercules, CA, USA) and the iTaq Universal SYBR Green One-Step Kit (BioRad, Hercules, CA, USA), respectively, following manufacturer’s instructions in 96-well plates (Applied Biosystems, Foster City, CA, USA). Primer sequences and annealing temperatures are indicated in Table 1. Gene expression was analyzed by CFX Manager™ Software (BioRad, Hercules, CA, USA) and Ct values were normalized against *D. rerio* glyceraldehyde-3-phosphate dehydrogenase (gapdh) gene.

### 2.6. Histopathology

Euthanized fishes were routinely processed for hematoxylin-eosin and Ziehl-Neelsen staining as was previously described [2,3]. The small size of the zebrafish permitted sagittal examination of each individual entirely in order to evaluate the presence of tuberculous granulomas with HE staining and acid-fast bacteria with ZN staining. All slides were independently examined by two blinded and experienced observers (M.A.R. and E.F.C.).

### 2.7. Immunohistochemistry

Immunohistochemistry was performed using the avidin-biotin-peroxidase complex (ABC) method to detect mycobacteria as described in Risalde et al. [10]. The primary antibody used was a rabbit polyclonal antiserum against *M. bovis* (Dako, Glostrup, Denmark) at 1:4000 dilution and the secondary antibody used was a biotinylated goat anti-rabbit IgG (Vector Laboratories, Burlin-game, CA, USA) at 1:2000 dilution. The pretreatment for antigen retrieval was incubation with 0.2% proteinase K (Sigma-Aldrich, Misuri, USA) in Tris buffer at 37°C for 8 min.

Tissues from zebrafish previously infected with *M. marinum* were included as positive controls [10]. Mycobacteria were identified based on positive immunolabeling by two experienced observers (M.A.R. and E.F.C.) and samples were classified as *Mycobacterium* positive or negative.

### 2.8. Statistical analysis

Data were assessed to calculate mean ± standard deviation (SD) or error (SE) values and were analyzed with the SPSS statistical software package (V.24.0; IBM, Somers, New York, USA). As data were normally distributed, differences in mycobacteria burden, immunological mediators and pathological changes between groups were tested through ANOVA and post-hoc test. *P* values ≤ 0.05 were considered statistically significant.

## 3. Results

### 3.1. *M. marinum* burden

Mycobacterial infection was confirmed by amplification of *M. marinum* 16S rDNA in zefrabish intestine, and by immunolabeling of *Mycobcterium* proteins and Ziehl-Neelsen staining in the whole fish.

*M. marinum* DNA levels tended to be lower in the groups immunized with HIMB when compared with groups immunized with iBCG for both administration routes of the immunostimulant (Figure 2A). Similarly, the number of fishes presenting positive immunolabelled cells tended to be lower in the group immunized IP with HIMB compared with the group immunized with iBCG (Figure 2B), but it was similar in both groups immunized by immersion. However, none of the differences resulted statistically significant. Of note, no acid-fast bacteria were detected.

**Figure 2.**
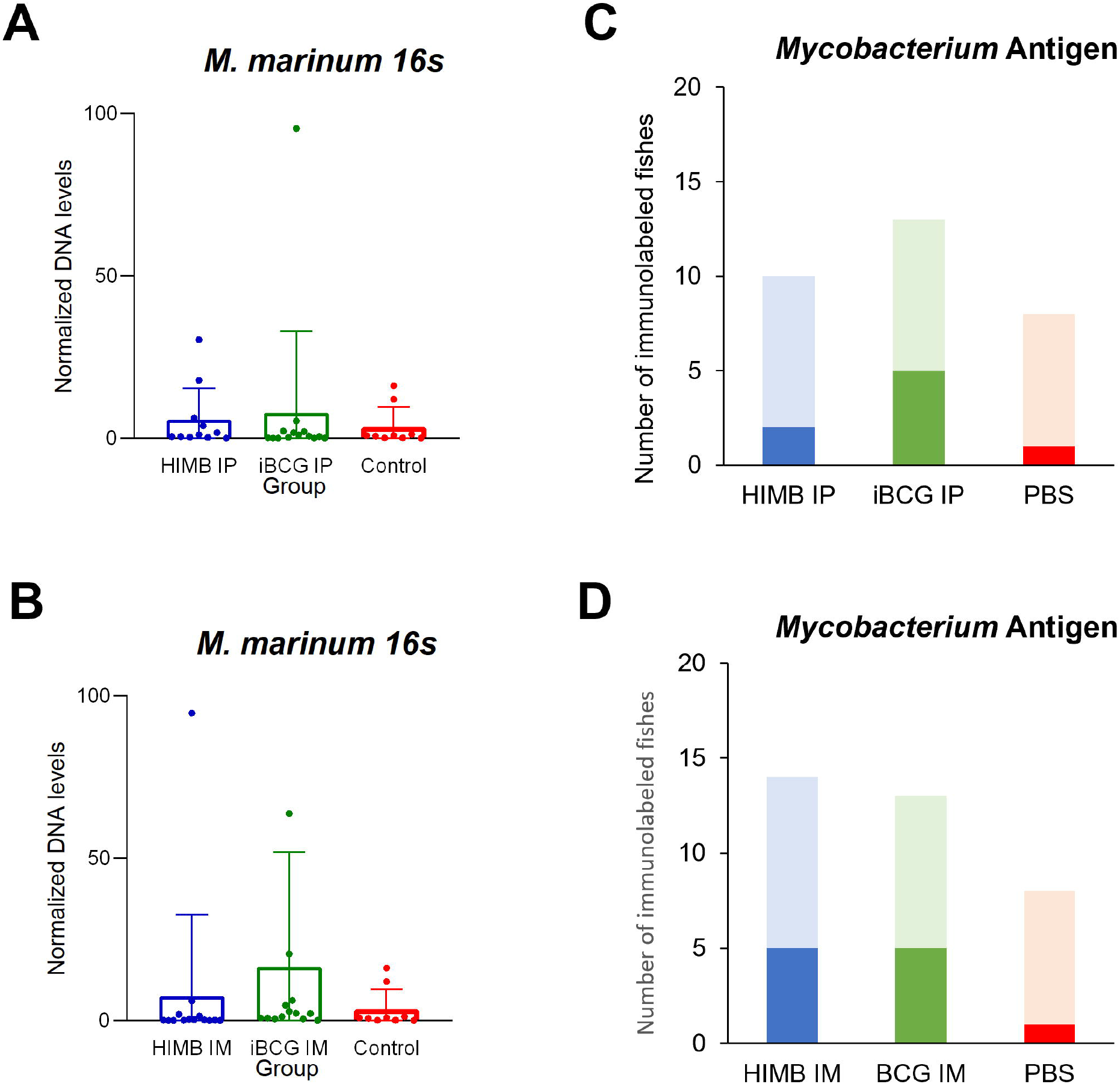
*Mycobacterium marinum* burden. Mean ± SD of normalized gene expression levels of *M. marinum* 16s in fishes immunized via intraperitoneal and controls (A), and via immersion and controls (B) at the end of the experiment. Number *Mycobacterium* immunolabeled fishes immunized via intraperitoneal and controls (C), and via immersion and controls (D) at the end of the experiment.

### 3.2. Microscopic findings

The presence of TB compatible lesions, such as granulomas, was evaluated by HE staining in the whole fish. No granulomas were observed in any of the groups.

### 3.3. Immune response

Relative expression of genes involved in innate immunity was measured in the intestine to assess the effect of the immunostimulant and its administration route on the immune response (Figures 3 and 4). Among all cytokines and receptors evaluated, Toll-like receptor (TLR) 4 levels were the highest in all groups. Gene expression level tended to be higher in the group treated with IP iBCG, although the difference was only significant for TLR2 and TLR4 between groups IP immunized (Figure 3; *p* < 0.0001).

**Figure 3.**
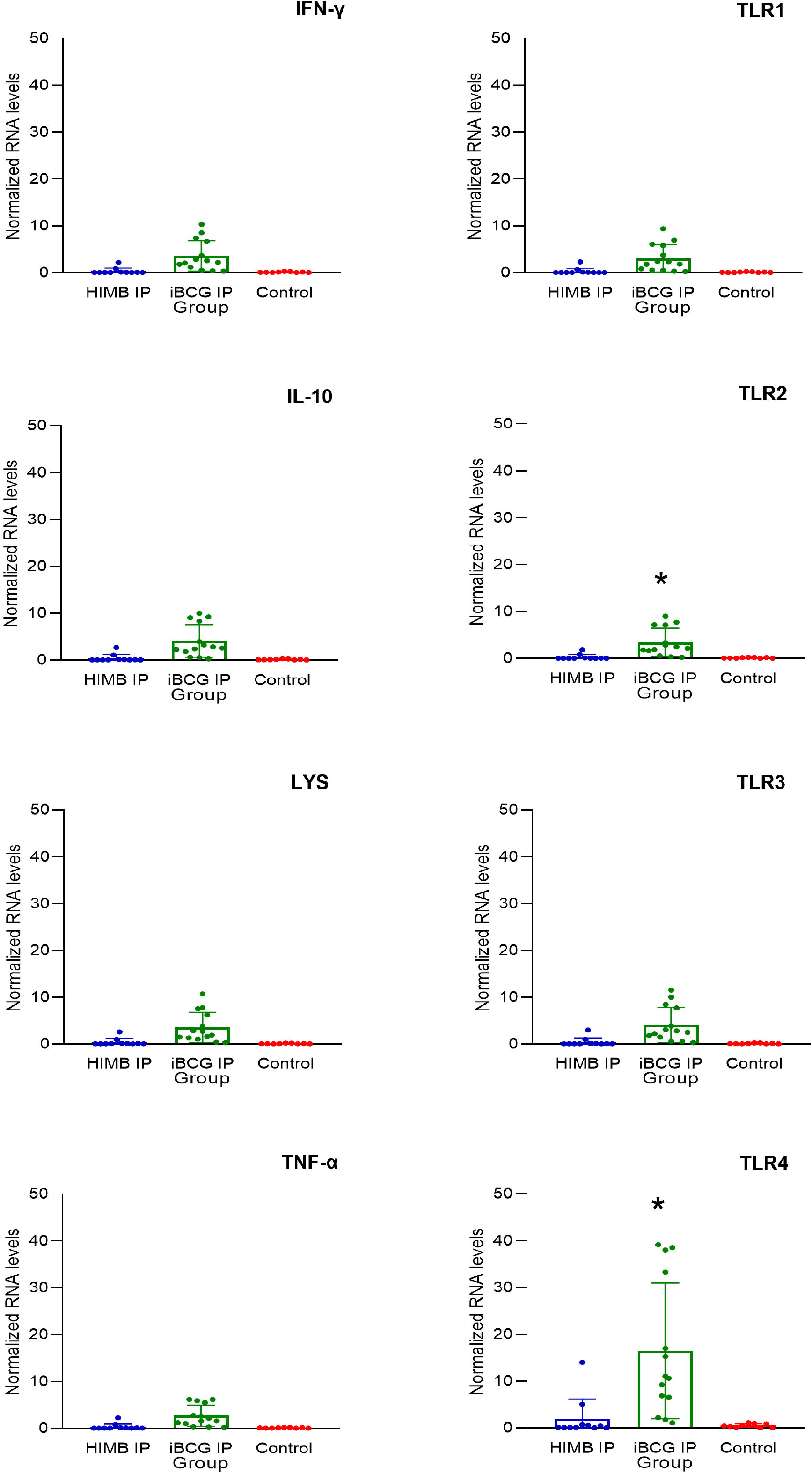
Immune mediators in intraperitoneally immunized fishes. Mean ± SD of normalized gene expression levels of interferon gamma (IFN-γ), interleukin 10 (IL-10), lysozyme (LYS), tumoral necrosis factor alpha (TNF-α), Toll-like receptor (TLR) 1, 2, 3, 4 in fishes immunized via intraperitoneal and controls at the end of the experiment.

**Figure 4.**
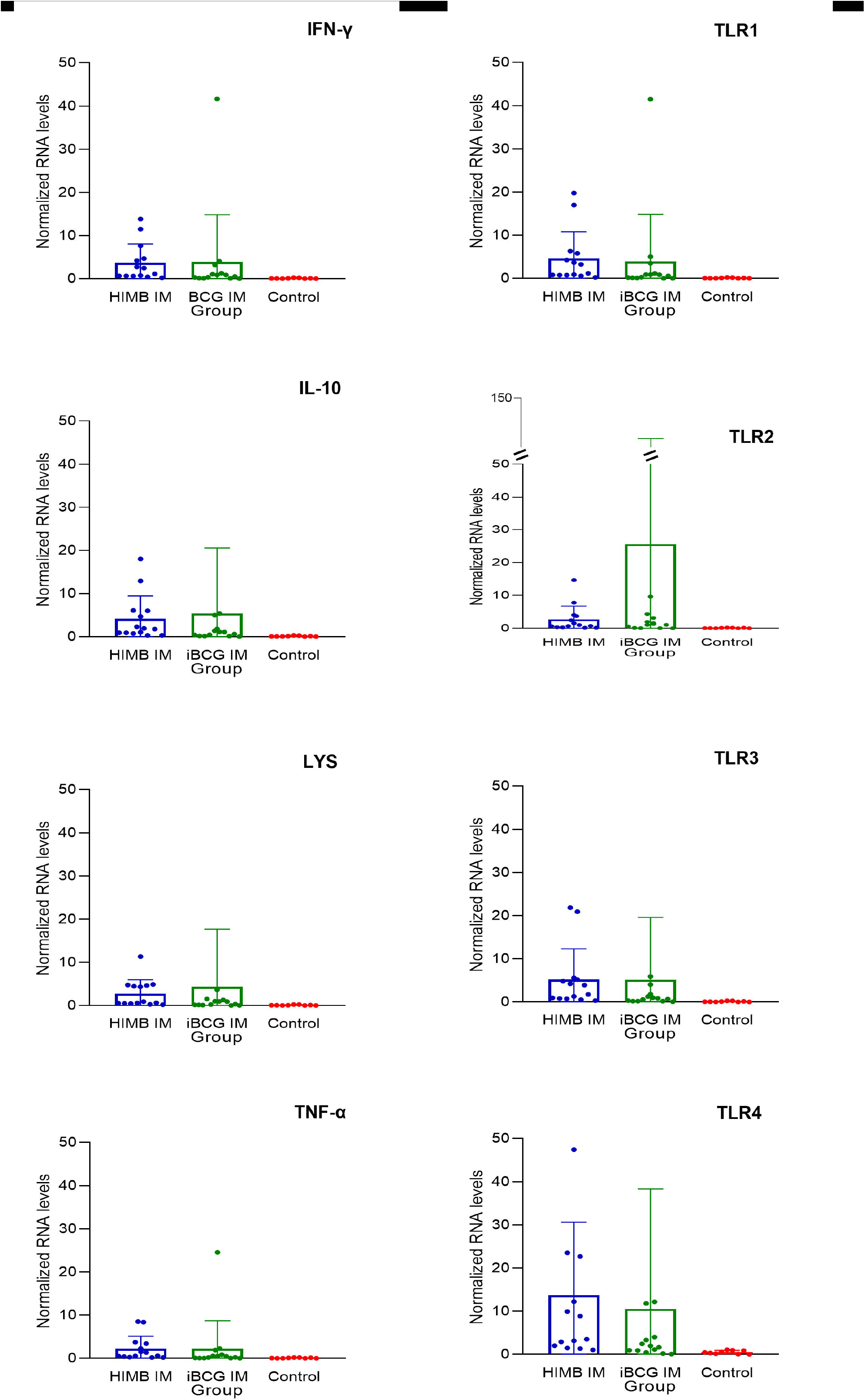
Immune mediators in fishes immunized by immersion. Mean ± SD of normalized gene expression levels of interferon gamma (IFN-γ), interleukin 10 (IL-10), lysozyme (LYS), tumoral necrosis factor alpha (TNF-α), Toll-like receptor (TLR) 1, 2, 3, 4 in fishes immunized via immersion and controls at the end of the experiment.

## 4. Discussion

Previous results support the use of vaccines containing inactivated mycobacteria to control fish mycobacteriosis [4]. Based on these evidences, herein we explored the performance of two inactivated *M. bovis*, HIMB and iBCG, against sublethal mycobacteriosis in the zebrafish model of tuberculosis mimicking natural conditions. In the present study, zebrafish were then infected with a sublethal dose of *M. marinum* via immersion. Since the most common infection routes for fish pathogens are the mucosa of the gills, skin and gut [1], challenge by immersion constitutes a more realistic approach than intraperitoneal injection [2,3]. However, immersion challenge implies certain heterogeneity in the infection dose received by each individual. In addition, a sublethal infection requires a low infection dose. Unlike previous studies that demonstrated a strong reduction in mycobacteria burden in fishes immunized with HIMB [2,3], the infection conditions in our study may have led to a lack of differences in mycobacteria levels between the immunized and control groups. Furthermore, due to unspecific mortality during the adaptation period, the number of fishes was not homogenous between groups, so the total *M. marinum* concentration in each tank, which was calculated according to the group size, varied among groups.

Despite the lack of effect of both immunostimulants (HIMB or iBCG) in mycobacterial load under the conditions of the present study, fishes parenterally immunized with iBCG showed a notorious increase in TLR2 and TLR4 gene expression. Both TLR2 and TLR4 are considered major pattern recognition receptors (PRRs) in the host defense against infection with Mycobacterium spp. [8,11]. In particular, Hu et al. [8,12] demonstrated that mutant zebrafish lacking TLR2 are more susceptible to infection with *M. marinum* due to impaired macrophage function. Accordingly, previous studies suggested the activation of TLR signaling pathways as the protective mechanism of HIMB [2,3] with possible implications in trained immunity [6]. Nevertheless, whether the protection level against fish mycobacteriosis differs between the two inactivated *M. bovis* strains used in the present study (HIMB or iBCG) remains unknown, especially when administered via immersion.

In conclusion, the results of our study demonstrated that the TLR2 and the TLR4 signaling pathways are involved in the immune response elicited by inactivated mycobacteria in the zebrafish model of tuberculosis. Overall, our findings support the use of inactivated mycobacteria in vaccine formulations for the control of mycobacteriosis. The development of mucosal vaccination strategies against pathogens is a priority in aquaculture [13], and our results encourage further research to standardize the mucosal route for routinary vaccination programs in fishes.

