## Supplementary material for "Heat-inactivated mycobacteria activate the Toll-like receptor 2 and 4 pathways in the zebrafish model of tuberculosis": Table

Table 1. Oligonucleotide primer sequences and annealing conditions.

| Gene | Forward primer | Reverse primer | Annealing conditions | Reference |
|---|---|---|---|---|
| GADPH | 5’-CGTGGTGCCAGTCAGAACAT-3’ | 5’-AGTCAGTGGACACAACCTGG-3’ | 56 ºC, 30 sec | Pacheco et al. (2020) |
| IFN-γ | 5’-GAGAGGCTGGCACATGTTCA-3’ | 5’-CCCATAGCGTTTCTGCATACG-3’ | 60 ºC, 1 min | Faikoh et al. (2014) |
| IL-10 | 5’-TCACGTCATGAACGAGATCC-3’ | 5’-CCTCTTGCATTTCACCATATCC-3’ | 60 ºC, 1 min | Faikoh et al. (2014) |
| LYS | 5’-CGTGGATGTCCTCGTGTGAAG-3’ | 5’-CCAATGGAGAATCCCTCAAA-3’ | 60 ºC, 1 min | Faikoh et al. (2014) |
| TNF-α | 5’-GCTTATGAGCCATGCAGTGA-3’ | 5’-TGCCCAGTCTGTCTCCTTCT-3’ | 56 ºC, 30 sec | Pacheco et al. (2020) |
| TLR1 | 5’-CAGAGCGAATGGTGCCACTAT-3’ | 5’-GTGGCAGAGGCTCCAGAAGA-3’ | 60 ºC, 1 min | Faikoh et al. (2014) |
| TLR2 | 5’-TGAATGGGTCGAGGAGATTC-3’ | 5’-CACAAAGTGCTCCGACAGAA-3’ | 56 ºC, 30 sec | Pacheco et al. (2020) |
| TLR3 | 5’-TGGAGCATCACAGGGATAAAGA-3’ | 5’-TGATGCCCATGCCTGTAAGA-3’ | 60 ºC, 1 min | Faikoh et al. (2014) |
| TLR4 | 5’-TCACCTGGACAGCAAGAATG-3’ | 5’-CGATTGACTTCCCTGCTTGA-3’ | 56 ºC, 30 sec | Pacheco et al. (2020) |

GADPH: glyceraldehyde-3-phosphate dehydrogenase; IFN-γ: interferon gamma; IL-10: interleukin 10; LYS: lysozyme; TNF-α: Tumor necrosis factor alpha; TLR: Toll-like receptor
